# Transcriptional plasticity drives leukemia immune escape

**DOI:** 10.1101/2021.08.23.457351

**Authors:** Kenneth Eagle, Taku Harada, Jérémie Kalfon, Monika W. Perez, Yaser Heshmati, Jazmin Ewers, Jošt Vrabič Koren, Joshua M. Dempster, Guillaume Kugener, Vikram R. Paralkar, Charles Y. Lin, Neekesh V. Dharia, Kimberly Stegmaier, Stuart H. Orkin, Maxim Pimkin

**Author notes:** These authors contributed equally (Kenneth Eagle, Taku Harada). These authors jointly supervised (Stuart H. Orkin, Maxim Pimkin).

## Abstract

Relapse of acute myeloid leukemia (AML) after allogeneic bone marrow transplantation (alloSCT) has been linked to immune evasion due to reduced expression of major histocompatibility complex class II (MHC-II) proteins through unknown mechanisms. We developed CORENODE, a computational algorithm for genome-wide transcription network decomposition, that identified the transcription factors (TFs) IRF8 and MEF2C as positive regulators and MYB and MEIS1 as negative regulators of MHC-II expression in AML cells. We show that reduced MHC-II expression at relapse is transcriptionally driven by combinatorial changes in the levels of these TFs, acting both independently and through the MHC-II coactivator CIITA. Beyond the MHC-II genes, MYB and IRF8 antagonistically regulate a broad genetic program responsible for cytokine signaling and T-cell stimulation that displays reduced expression at relapse. A small number of cells with altered TF levels and silenced MHC-II expression are present at the time of initial leukemia diagnosis, likely contributing to eventual relapse. Our findings reveal an adaptive transcriptional mechanism of AML evolution after allogenic transplantation whereby combinatorial fluctuations of TF levels under immune pressure result in selection of cells with a silenced T-cell stimulation program.

## INTRODUCTION

Treatment of acute myeloid leukemia (AML) with curative intent often utilizes allogeneic hematopoietic stem cell transplantation (alloSCT), which relies on the graft-versus-leukemia effect (GvL), whereby donor-derived T-cells eliminate residual leukemia cells (*1*). While the immunologic mechanisms of GvL are complex, relapse of AML after allogeneic transplantation has been linked to the loss of major histocompatibility complex class II (MHC-II) expression (*2–4*). Notably, treatment of AML blasts with IFNγ restored MHC-II expression, and no new coding DNA mutations were detected in comparative analyses of pre- and post-alloSCT relapsed samples (*3, 4*). Thus, the reduced expression of MHC-II at relapse appears to have a regulatory basis. Decreased levels of the transcriptional coactivator CIITA were observed in some cases (*4*), but loss of MHC-II expression was also seen despite unchanged or elevated levels of CIITA (*3*), suggesting alternative mechanisms of immune escape.

## RESULTS

### Network decomposition by combinatorial regression

To identify the transcriptional regulators of MHC-II in AML, we sought to reverse engineer a transcription regulatory network from public genome-scale datasets (Fig. 1a). In principle, the expression level of a gene in a given context may be computationally inferred from the identities of its transcriptional regulators, their cellular levels and the gene-regulatory function that represents the quantitative relationship between a transcription factor (TF) and its target gene (*5, 6*). We began by defining a list of core regulatory (CR) TFs for AML lineage specification (*7*). CR TFs are typically marked by extended, closely spaced enhancers with markedly high levels of histone acetylation and cofactor recruitment, termed super-enhancers (SEs) (*8*). Among a chromatin immunoprecipitation sequencing (ChIP-seq) dataset for the enhancer histone mark H3K27ac in a panel of 49 primary human AML samples (*9*), we identified 1298 SEs present in at least 10 (~20%) samples (Supplementary Fig. 1). These common SEs were proximal to 2748 genes, of which 220 encoded TFs. Since CR TFs are typically essential for lineage survival (*10, 11*), we focused this list further by asking which of the 220 SE-associated TFs were dependencies selective for AML versus other malignancies. We interrogated data from the Broad Cancer Dependency Map project, a collection of genome-scale CRISPR knockout screens of 18,119 genes in 769 cell lines, including 19 AML lines (*12, 13*). Applying a skewed-LRT test (*14*) to compare guide RNA drop out between AML and non-AML cell lines, we identified 40 TFs selectively essential in AML (Supplementary Fig. 1). We defined the AML core regulatory circuit (CRC) by intersecting the 40 essential TFs with the 220 SE-driven TFs, resulting in 19 CR TFs (Fig. 1a).

**Fig. 1.**
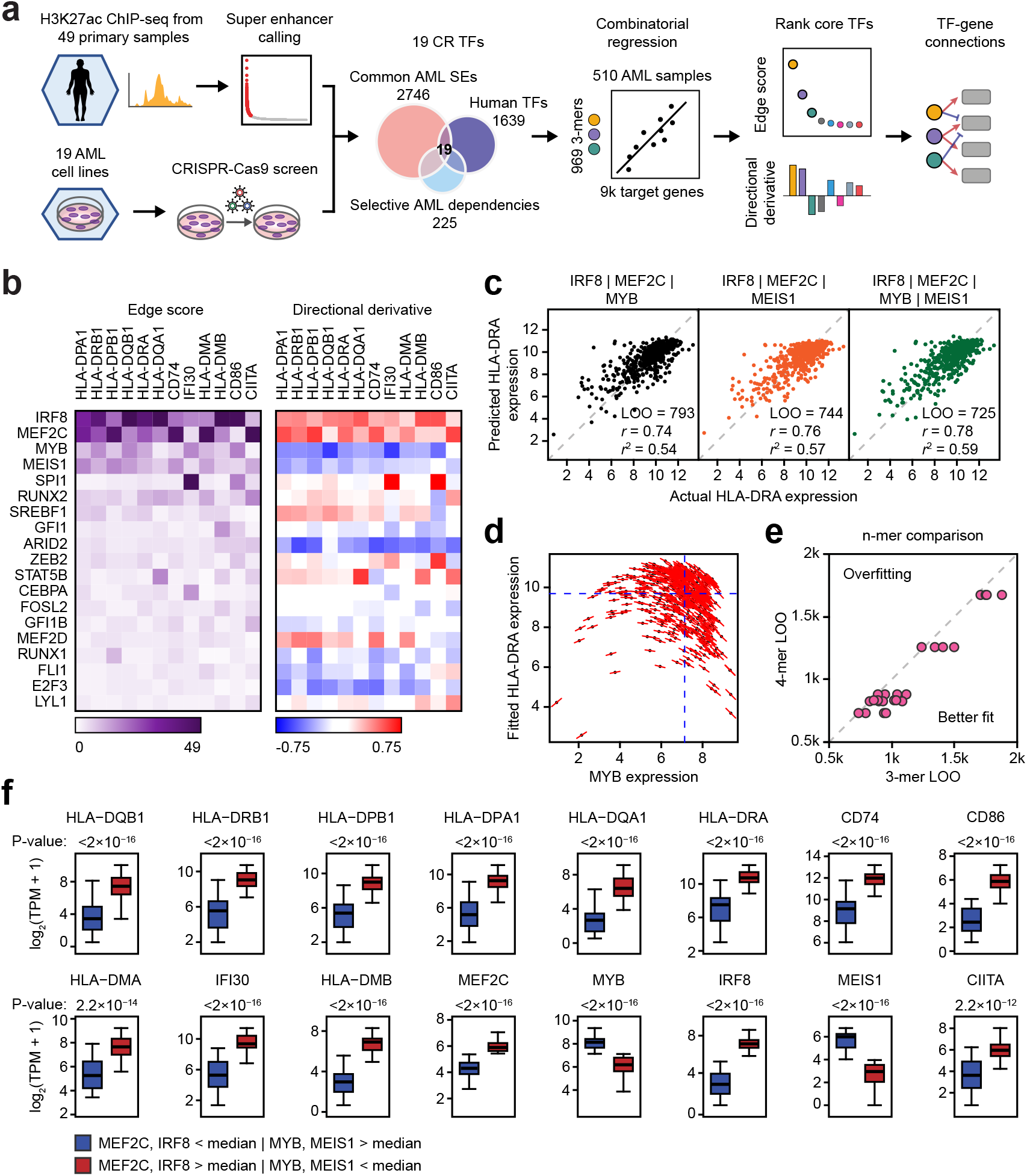
CORENODE identifies a tetrad of TFs regulating MHC-II genes in AML. **a**, Study schematic. **b**, Heatmaps of edge scores (ES) and directional derivatives (DD) representing computationally established edges between target genes (columns) and TFs (rows, sorted by average ES). Higher ES corresponds to higher confidence edges, while DD predicts amplitude and directionality (positive *vs*. negative) of TF-target regulation. **c**, 3- and 4-mer CORENODE fits for the gene HLA-DRA with indicated goodness-of-fit metrics. **d**, A graphic illustration of DD. MYB expression is plotted against the CORENODE fitted HLA-DRA expression using a IRF8/MEF2C/MYB 3-mer. Red sticks indicate MYB slopes (aggregated from the 4 MYB-containing terms: linear, quadratic and two cross terms) in each sample. Aggregation of all MYB slopes for all samples produces MYB DD. **e**, LOO improvement between 3-mers (4 per gene [4 combinations of 4 TFs taken 3 at a time]) and the 4-mer (1 per gene, combining all 4 TFs) for 6 MHC-II genes. **f**, Gene expression in two populations of the BeatAML dataset (*15*). The blue population (*n*=46) represents patient samples with below-median expression of IRF8 and MEF2C and above-median expression of MYB and MEIS1, while the red population (*n*=56) represents patient samples with the opposite pattern of TF expression.

We sought next to define the gene-regulatory functions linking CR TFs to all expressed genes. We developed CORENODE (COmbinatorial REgression for NetwOrk DEcomposition), a coexpression-based network decomposition algorithm where TF-target gene connections, or edges, are inferred from naturally occurring variation in gene expression within a reasonably large set of samples. We hypothesized that the expression level of each gene could be estimated from the expression levels of a small subset, possibly 3-5 TFs, of the entire CR circuit. Given that the intersection of dependency and SE profiling data nominated only 19 TFs, it was computationally feasible to design a comprehensive combinatorial regression approach to fit expression of each target gene against all possible combinations of the CR TFs (Fig. 1a, Supplementary Fig. 1). Using the protein-coding transcriptomes of 510 primary AML samples from the BeatAML study (*15*), we regressed the mRNA levels of all genes in the expressed genome against mRNA levels of the 19 CR TFs taken 3 at a time, using linear, quadratic, and cross terms. Use of 3-mers provided the best balance between fit quality and overfitting risk genome-wide. Thus, for every target gene we generated 969 gene-regulatory functions corresponding to all possible 3-mer combinations of the 19 TFs. We chose the top 5% of best-fitting 3-mers and calculated an edge score (*ES*) for each TF-target gene pair as the number of times the TF appeared in the top 5% of best fits. Finally, we estimated the direction and amplitude of positive or negative regulation by aggregating response slopes for each TF across all samples and 3-mers, which was termed directional derivative (*DD*).

### A TF tetrad regulates MHC-II genes in AML

CORENODE prioritized four TFs (IRF8, MYB, MEF2C and MEIS1) as candidate regulators of the MHC-II and several additional genes involved in antigen presentation that have been associated with immune escape in AML (CD74, IFI30, HLA-DMA, HLA-DMB, CD86) (*3*) (Fig. 1b). MYB and MEIS1 were predicted to negatively regulate expression of these genes, while IRF8 and MEF2C were predicted to be positive regulators (Fig. 1b-d). Combining these TFs in a single 4-mer resulted in a modest improvement of the regression fits (compared to 3-mers) without overfitting (Fig. 1c, Supplementary Fig. 2).

We examined how variation in the levels of the TF tetrad correlated with changes in MHC-II expression in two independent datasets, BeatAML and TCGA, including 510 and 151 patients, respectively (*15, 16*). In both datasets the expression of MHC-II and other genes associated with immune escape in patients with high (above the median) IRF8/MEF2C and low (below the median) MYB/MEIS1 was up to 17-fold higher than in patients with the opposite pattern of TF expression (Fig. 1f, Supplementary Fig. 2). To validate the impact of these TFs on MHC-II regulation, we inactivated each TF with CRISPR/Cas9 editing followed by RNA-seq in an AML cell line (Supplementary Fig. 3). Consistent with predictions of CORENODE, loss of IRF8 and MEF2C led to reduced expression of MHC-II and other genes associated with immune escape, while loss of MYB and MEIS1 was associated with increased expression (Fig. 2a). We confirmed these findings at the protein level by measuring cell surface expression of MHC-II molecules after TF knockout in a panel of AML cell lines (Fig. 2b, Supplementary Fig. 3). Overall, MYB and IRF8 had a major impact on MHC-II expression, while the effects of MEIS1 and MEF2C were subtle and less consistent.

**Fig. 2.**
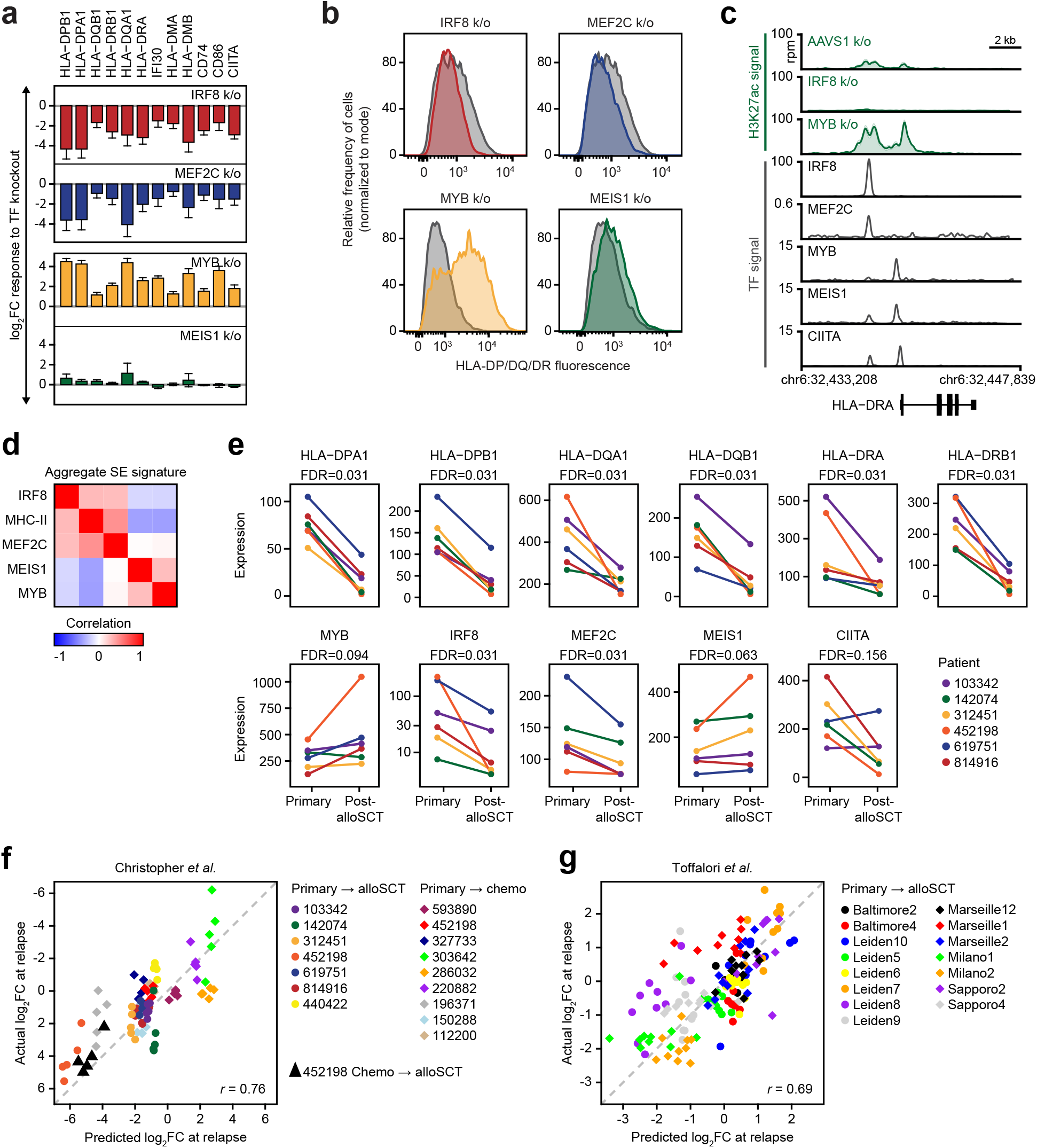
IRF8, MEF2C, MYB and MEIS1 regulate MHC-II expression in AML. **a**, Transcriptional response of the indicated genes measured by mRNA-seq 72 hours after TF knockout. Error bars represent standard error. MEF2C knockout changes were not significant after genome-wide adjustment for multiple hypothesis testing. Among MEIS1 knockout-induced changes only HLA-DRA and HLA-DQB1 had *q*-values <0.1. **b**, Changes in the surface expression of the MHC-II molecules detected by immunostaining with a pan-specific anti-HLA-DP/DQ/DR antibody following TF knockout. IRF8 and MEF2C knockouts were evaluated at 96 hrs post-electroporation. MYB and MEIS1 knockouts were evaluated at 72 hrs. An AAVS1 (“safe harbor”) targeting sgRNA was used as a control. **c**, H3K27 acetylation and TF binding in the HLA-DRA locus. Green tracks are H3K27ac metatracks composed by overlaying semi-transparent area plots representing 2 biological replicates of the indicated TF knockouts compared to AAVS1 control. The average profile is represented by a thick line. Grey tracks represent binding of the indicated TFs. All of the shown ChIP-seq experiments were performed in MV411 except for CIITA which was downloaded from Scharer *et al*. (*29*) and represents a lymphoid cell line. Refer to Supplementary Fig. 4 for a map of H3K27 acetylation and TF binding in the entire MHC-II locus and genome-wide analysis of H3K27ac changes. **d**, Similarity matrix of SE scores associated with the TF tetrad and MHC-II genes using data from McKeown *et al*. (*9*). **e**, mRNA expression in 6 paired primary and post-alloSCT samples from Christopher *et al*. (*3*) **f,g**, CORENODE accurately predicts MHC-II expression changes at relapse. Predicted by *vs*. observed log_2_ fold change in MHC-II expression in paired samples from Christopher *et al*. (*3*) (**f**) and Toffalori *et al*. (*4*) (**g**) between initial presentation and relapse. For each patient (color coded as indicated) each datapoint reflects predicted and observed changes in the expression of one MHC-II gene. The IRF8/MYB/MEIS1/MEF2C 4-mer was used in panel **f**, while CIITA was added as a 5^th^ term in panel **g** resulting in a better fit.

The MHC-II locus contains several SEs whose H3K27 acetylation levels correlate with expression of MHC-II genes in primary AML cells (Supplementary Fig. 4). Spike-in controlled ChIP-seq analysis detected a significant increase in H3K27 acetylation at these SEs after MYB knockout, despite the expected global decrease of H3K27 acetylation elsewhere across the genome (Fig. 2c, Supplementary Fig. 4). In contrast, IRF8 knockout was accompanied by loss of H3K27 acetylation. Furthermore, the SEs associated with IRF8 and MEF2C in primary AML blast cells positively correlated with MHC-II SEs, while SEs controlling MYB and MEIS1 negatively correlated (Fig. 2d). These data provided further evidence of antagonistic functions of these TFs in MHC-II regulation.

We next examined how MHC-II gene expression after alloSCT correlated with expression of the TF tetrad. We accessed the published RNA-seq data from paired AML samples at initial presentation and relapse after alloSCT, where decreased MHC-II expression was observed in 6 of the 7 examined patients (*3*). Despite substantial variability of baseline expression, we observed significant down-regulation of IRF8 and, to a lesser extent, MEF2C in all patients with reduced MHC-II expression at relapse (Fig. 2e). In contrast, MYB and MEIS1 were upregulated in 5 of 6 patients at relapse, which was of borderline statistical significance. Each of these changes would be expected to lead to reduced expression of MHC-II genes.

### A combinatorial transcriptional mechanism of immune escape

To this point, our computational predictions and knockout experiments demonstrated that perturbation of any one of the 4 candidate TFs was sufficient to effect a change in MHC-II expression, while examination of AML relapses revealed changes in the levels of these TFs in the direction consistent with reduced MHC-II expression. However, the degree of observed TF change was often modest and, in the case of MYB and MEIS1, not evident in all post-alloSCT samples (Fig. 2e). This observation led us to hypothesize that immune escape may be enacted by combinatorial changes in the tetrad. To assess whether the observed changes in TF expression adequately accounted for the magnitude of MHC-II downregulation at the time of relapse, we used the CORENODE gene-regulatory functions to predict the change in MHC-II expression from the pre- and post-relapse expression values of the TF tetrad. For additional statistical robustness we analyzed all paired samples from the two available studies (*3, 4*), including patients regardless of therapy (chemotherapy *vs*. alloSCT) or the direction of MHC-II change. In both datasets, the predicted changes in MHC-II expression closely correlated with the actual changes observed at the time of relapse (Fig. 2f-g). Given that the gene-regulatory functions were derived from an unrelated dataset, these findings provided validation for the computational model and confirmed the combinatorial action of CR TFs in MHC-II regulation.

### MYB and MEIS1 act independently of CIITA

The MHC-II genes are regulated by CIITA, a transcriptional coactivator that does not bind directly to DNA and acts as a histone acetylase (*17–19*). Expression of MHC-II genes correlates with CIITA expression (Fig. 3a, Supplementary Fig. 5). Therefore, we asked if the TF tetrad regulates MHC-II expression via CIITA. Indeed, CORENODE identified MEF2C, MEIS1 and IRF8 as candidate regulators of CIITA (Fig. 1b). Knockouts of IRF8 and MEF2C led to decreased CIITA expression, whereas a MYB knockout was associated with increased expression (Fig. 2a). Thus, at least 3 of the 4 TFs appear to participate in MHC-II expression by modulating CIITA levels. However, post-alloSCT silencing of the MHC-II genes was not consistently associated with reduced CIITA expression (Fig. 2e), pointing to an independent mechanism of MHC-II regulation. In addition, ChIP-seq confirmed binding of the entire tetrad at the MHC-II locus (Figs. 2c, Supplementary Fig. 4), consistent with a possibility of direct action. These conflicting observations prompted us to investigate further CIITA dependent and independent actions of the tetrad. First, we repeated CORENODE regressions of MHC-II genes using 4-mers and 5-mers where CIITA was used as an additional regression term. Inclusion of CIITA in the model resulted in a marked improvement of goodness-of-fit for all genes in the MHC-II locus without overfitting (Fig. 3a,b). Furthermore, *t*- and *p*-values of individual regression terms, representing probability of their independent contribution to the overall fit, indicated that the TFs were independent variables, with the strongest contributions from MYB and CIITA (Fig. 3c). This analysis suggested that members of the TF tetrad—and particularly MYB— regulate MHC-II in a CIITA-independent manner. To confirm this inference, we asked if the negatively acting TFs regulate MHC-II expression in the absence of CIITA coactivation. We generated a CIITA knockout cell line (Supplementary Fig.5). As predicted, baseline expression of MHC-II proteins was reduced in CIITA-deficient cells (Fig. 3d). Nonetheless, loss of MYB and MEIS1 led to increased MHC-II expression (Fig. 3d,e), consistent with a CIITA-independent mechanism of MHC-II regulation by these TFs.

**Fig. 3.**
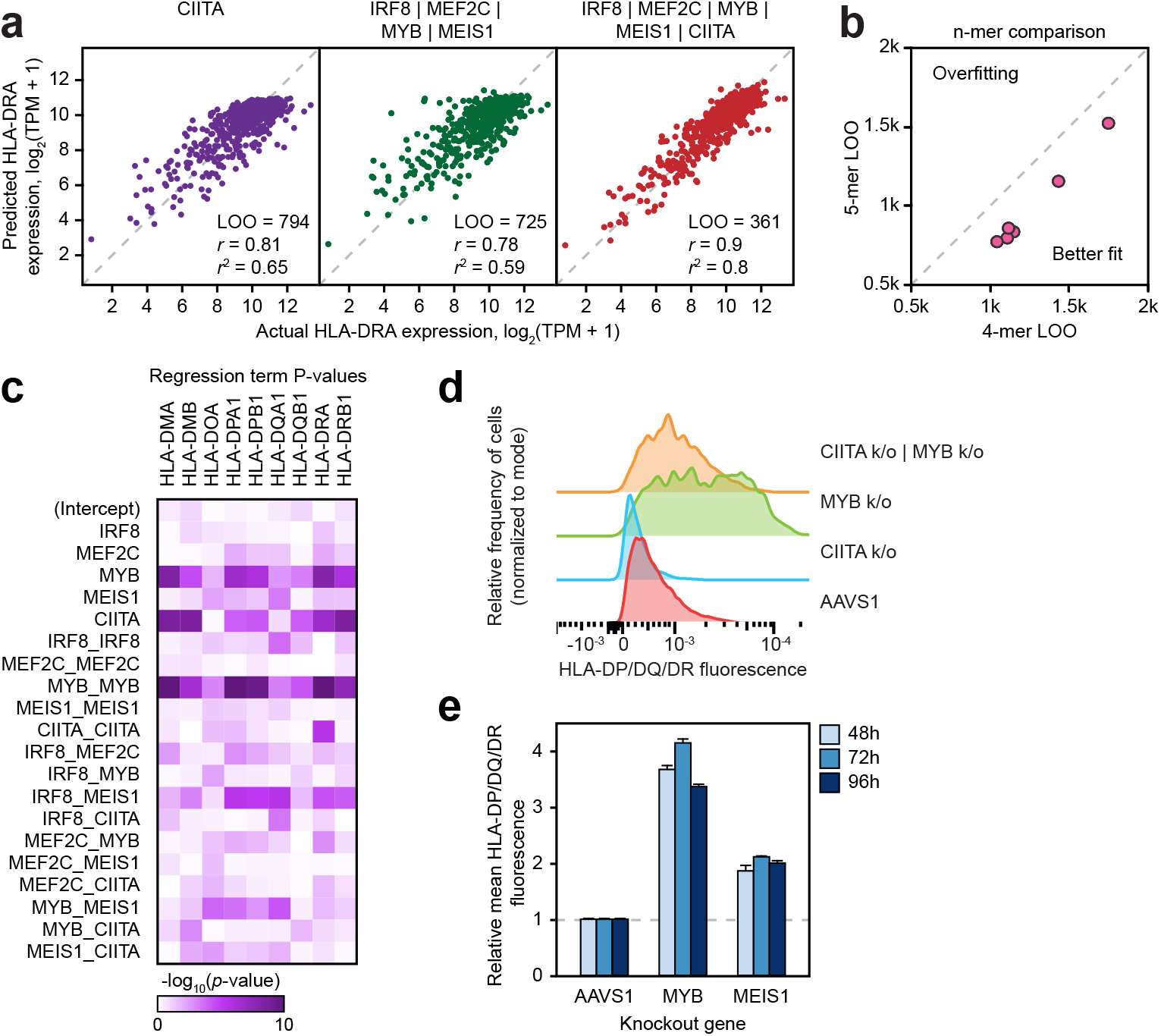
CIITA-dependent and independent regulation of MHC-II genes by the TF tetrad. **a**, 1-mer CORENODE fit using CIITA (linear + quadratic term) as the only predictor of HLA-DRA expression, as well as 4- and 5-mer CORENODE HLA-DRA fits with indicated goodness-of-fit metrics. **b**, LOO improvement between TF 4-mers (6 fits, 1 per HLA gene) and 5-mers that include CIITA in addition to the 4 TFs. **c**, Heatmap visualization of regression term *p*-values representing probability of the term’s *t*-value being zero. Lower *p*-values, visualized by denser color on the heatmap, reflect higher probability and magnitude of the term’s independent contribution to the overall fit. The statistics are calculated separately for each gene using the 5-mer composed of the TF tetrad and CIITA. **d**, Change in the surface expression of HLA-DP/DQ/DR following MYB knockout in wild type MV411 and following CIITA knockout, detected by immunostaining with a pan-specific anti-HLA-DR/DP/DQ antibody. **e**, Changes in the surface expression of HLA-DP/DQ/DR in a CIITA knockout clone after indicated TF knockouts.

### IRF8 and MYB antagonistically regulate a GvL response module

We sought to place MHC-II genes in the context of genome-wide transcriptional regulation by the 19 CR TFs. We began by evaluating the global accuracy of CORENODE in predicting transcriptional regulation. To this end, we generated a list of high-confidence targets of each CR TF by applying a minimal edge score cutoff of 15, and cross-plotted directional derivatives for each target gene *vs*. the actual response observed on RNA-seq after knockout of the corresponding TF in MV411 cells. A significant correlation was observed between predicted and actual changes in gene expression following TF loss (Fig. 4a). Having confirmed CORENODE’s precision in predicting genomic regulation, we built a map of high-confidence edges in the AML transcription regulatory network, revealing distinct functional modules of CR TFs and coregulated genes (Fig. 4b). The model highlighted several TFs (MYB, FLI1, STAT5B, CEBPA, IRF8, SPI1) as genome-wide regulators, while other TFs (GFI1, MEIS1, RUNX1, RUNX2, MEF2C, MEF2D) were projected to regulate more narrow transcriptional programs. In addition, the model predicted partially antagonistic functions for some TFs on pairwise comparison (Supplementary Fig.5). We focused attention on the antagonistic actions of MYB and IRF8. CORENODE highlighted a module of 58 genes predicted to be upregulated by IRF8 and downregulated by MYB (Fig. 4c). The predicted outcome conformed closely to the actual responses following TF knockouts (Fig. 4d). The module is highly enriched for genes participating in multiple aspects of alloreactivity (Fig. 4e, Supplementary Fig. 5), including response to IL-2 and IFNγ (MAP2K1, OAS1, IFI30, MX1, TNFRSF1B), T-cell stimulation (CD86, BTN3A1, TNFSF8, SH2B3), adhesion (ITGB7, TGFBI), inflammasome formation (NLRP1) and transcription regulation (NOD2, SPI1, MALT1, CBX7, MAFB).

**Fig. 4.**
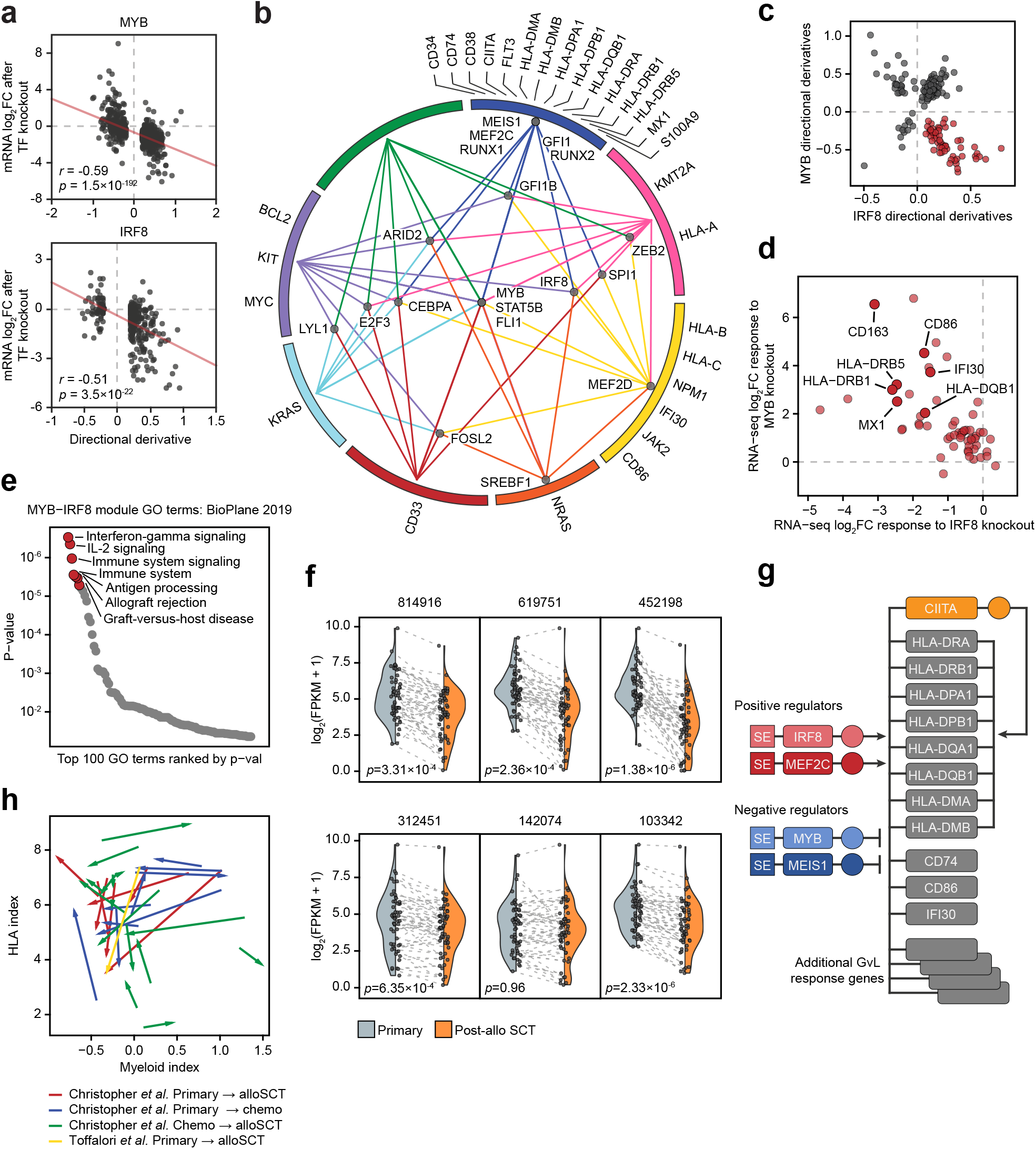
Global transcription network decomposition by CORENODE. **a**, CORENODE validation by TF knockout followed by mRNA-seq. DD values (predicted response) for high confidence MYB and IRF8 edges (ES ≥ 15 and [DD] ≥ 0.2) are plotted against actual log_2_ fold change in mRNA expression measured by RNA-seq 72 hours after MYB knockout (observed response). Spike-in control was used to account for the global transcriptional collapse after MYB knockout. **b**, A graphic representation of the AML transcription regulation network reconstructed from the CORENODE output. The circle represents clusters of target genes coregulated by the TFs, as illustrated by TF-cluster edges. Proximity of the TFs to the clusters reflects their specificity towards them (for example, MYB regulates all clusters while MEIS1 regulates only one cluster). A geometrically optimal map with the shortest aggregate distance of the TF-cluster edges was produced by solving for the unweighted Weber problem. **c**, CORENODE identifies a gene module predicted to be repressed by MYB and activated by IRF8. Visualized are DD scores of all genes with high confidence (ES ≥ 15) MYB and/or IRF8 edges; 58 of these genes (highlighted in red) are predicted to be negatively regulated by MYB and positively regulated by IRF8. **d**, Transcriptional responses of the 58 genes comprising the GvL module depicted in panel **c** to MYB and IRF8 knockouts measured by RNA-seq are cross-plotted, confirming the predicted directionality of regulation for the vast majority of genes. Spike-in normalization was not used for this plot. **e**, Gene set enrichment analysis of the MYB-IRF8 (GvL) module genes in panel **c** using Enrichr (*30*). **f**, Decreased expression of the GvL module in paired primary and post-alloSCT patient samples from Christopher *et al*. (*3*) with reduced MHC-II expression at relapse. **g**, Schematic of the proposed regulation of the MHC-II and other GvL genes in AML by the TF tetrad. **h**, MHC-II expression at relapse does not correlate with myeloid differentiation state. The index of myeloid differentiation is plotted against combined MHC-II expression in each paired sample and the direction of change between initial presentation and relapse is marked by arrows.

Importantly, the module was downregulated in 5 of 6 patients with reduced MHC-II expression after alloSCT, with the exception of the sole patient (142074) who displayed decreased MYB expression at relapse (Fig. 4f, Supplementary Fig. 5). Thus, antagonistic actions of MYB and IRF8 facilitate immune evasion by silencing a broad transcriptional program that regulates the interaction between AML blasts and donor T-cells (Fig. 4g).

### Immune escape is uncoupled from the myeloid differentiation state

Members of the TF tetrad have previously been implicated in the regulation of normal and malignant myelopoiesis: MYB and MEIS1 restrain myeloid differentiation while IRF8 and MEF2C promote it (*20–23*). Additionally, AML cells form a functional hierarchy of differentiation states arising from a leukemia stem cell (LSC) (*24*), and activation of immune pathways is a hallmark of monocytic development (*25*). Thus, we hypothesized that reduced expression of IRF8 and MEF2C and increased expression of MYB and MEIS1 at relapse reflect evolution of leukemia into a less differentiated state. To test this hypothesis, we computed a myeloid differentiation index from the mRNA expression of 19 cell type-specific markers, which correctly reproduced both the normal myeloid trajectory and the functional AML hierarchy (Supplementary Fig. 6). However, MHC-II expression correlated poorly with myeloid differentiation in the BeatAML and TCGA datasets, and changes in the MHC-II expression at relapse could not be predicted from the changes in the myeloid differentiation index (Fig. 4h and Supplementary Fig. 6). We conclude that reduced MHC-II expression at relapse is regulated independently from the myeloid differentiation state.

### Transcriptional plasticity underlies relapse

We examined expression of MHC-II genes in AML cells at the single-cell level. We analyzed single-cell transcriptomes obtained from patient 452198 at presentation and post-alloSCT relapse (*3*). Plotting all cells together using t-distributed stochastic neighbor embedding (t-SNE) revealed distinct clusters of AML cells corresponding to initial presentation and relapse (Fig. 5a). Although most AML cells displayed high MHC-II expression at presentation, approximately 1 in 40 cells had low or undetectable MHC-II levels similar to those seen at relapse (Fig. 5b,c). Some of these MHC-II-low cells co-clustered with the relapsed cell population, indicating similar global transcription patterns (Fig. 5c). The MHC-II-low cells displayed significantly higher MYB and lower IRF8 levels compared to the cells expressing high MHC-II levels at presentation (Fig. 5d). The average MYB levels in the MHC-II-low cell population at presentation were still lower, and the IRF8 levels were still higher than the average levels of these TFs at relapse. However, up to 1 in 6 cells in the MHC-II-low population (approximately 1 in 250 of all AML cells at presentation) expressed undetectable IRF8 and markedly elevated MYB levels that matched or exceeded the average MYB levels seen at relapse (Fig. 5c). Although expression of MEIS1 was increased and expression of MEF2C was decreased in most cells at relapse, we found no statistically significant change in their expression in the MHC-II-low cells at presentation, indicating that stable changes in these TFs were acquired later in the course of leukemia evolution (Fig. 5d). We conclude that a minor population of cells with reduced MHC-II expression driven by altered levels of MYB and IRF8 was present in this patient at the time of initial presentation and likely contributed to the subsequent relapse.

**Fig. 5.**
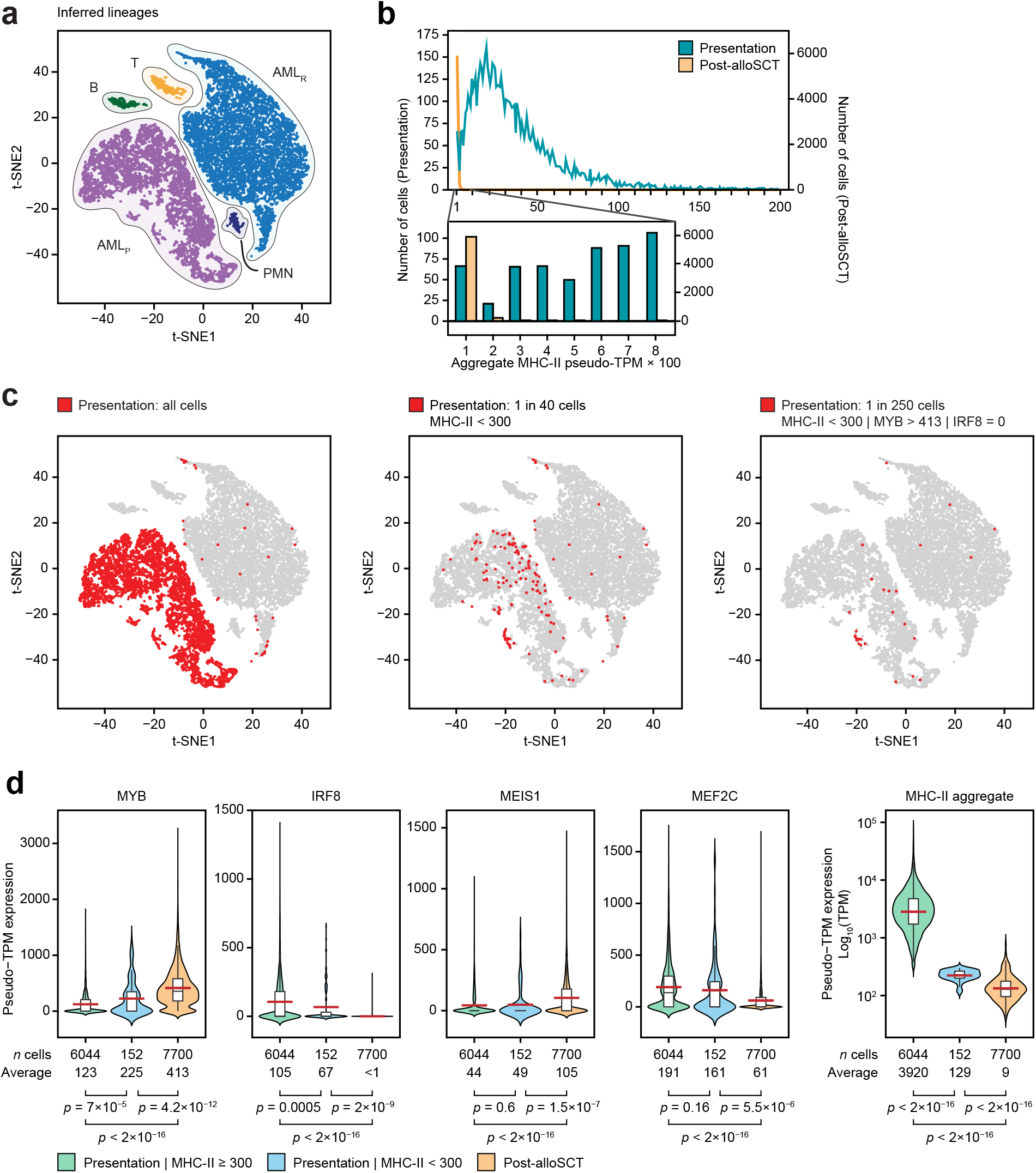
Single cell RNA-seq reveals an adaptive mechanism of transcriptional evolution underlying relapse. **a**, scRNA-seq of AML cells obtained at initial presentation (AML_P_) and post-alloSCT relapse (AML_R_). Data from patient 452198 were downloaded from Christopher *et al*. (*3*) and clustered according to their genome-wide expression patterns using t-distributed stochastic neighbor embedding (t-SNE). Cell lineages were inferred from expression of lineage markers. **b**, Distribution of MHC-II expression in AML cells. Individual cells are placed in bins according to their aggregate MHC-II expression and cell frequency is plotted for each bin. **c**, t-SNE plots start with the same plot as in panel **a** but highlight cells from the initial presentation that fit the designated criteria. **d**, Single-cell expression of the TF tetrad in AML cells with low and high aggregate MHC-II expression at presentation compared to relapse.

## DISCUSSION

By developing a network decomposition approach that captures combinatorial TF interactions, we identified a tetrad of core TFs that regulate immune escape of AML cells after allogeneic bone marrow transplantation. The tetrad TFs appear to act as opposing pairs: IRF8 and MEF2C positively control expression of MHC-II genes, whereas MYB and MEIS1 serve to downregulate them. IRF8 is an established effector of IFNγ and controls MHC-II expression in myeloid cells (*26, 27*). Remarkably, the transcriptional activator MYB acts as a strong negative regulator of MHC-II genes. Consistent with our observations, *Myb*-deficient mouse B-cells exhibit elevated MHC-II expression (*28*). Although MYB negatively regulates the MHC-II co-activator CIITA, both MYB and MEIS1 appear to function primarily in a CIITA-independent manner. Beyond reducing MHC-II expression, antagonistic actions of CR TFs facilitate immune escape by enacting a broader program of transcriptional changes resulting in reduced expression of multiple genes involved in antigen presentation, cytokine response and T-cell co-stimulation. Notably, to the extent that these core TFs are essential for AML survival, post-SCT relapse appears to rely on relatively modest changes in their expression that cooperatively mediate immune evasion without affecting cell viability. A small population of such cells with reduced MHC-II expression due to altered TF levels is present at the time of initial diagnosis and likely contributes to relapse under *in vivo* selection. Whether transcriptional heterogeneity is hard-wired (for example, by mutations in non-coding regulatory DNA elements) or driven by stochastic TF fluctuations remains to be elucidated. Nonetheless, we speculate that the immune pressure imposed by allogenic transplantation selects AML cells with a pattern of CR TF expression that provides an optimal balance between immune escape and overall fitness. Such “transcriptional evolution” by selection of cells with favorable transcriptional states mediated by altered combinatorial patterns of key TFs may drive progression of other tumors and eventual relapse.

## SUPPLEMENTARY FIGURE LEGENDS

**Supplementary Fig. 1. CORENODE development. a**, Data and computational flowchart. **b**, SE signal and ranks in a representative primary patient sample with high MHC type II expression. **c**, SE presence in 49 primary AML samples. Data in **c** and **d** are from Ref.(*9*). **d**, Transcription factors identified as selective AML dependencies by the Broad Cancer Dependency Map project. The heatmap reflects probability of dependency > 0.5. 40 TF genes with dependency probability >0.5 in 3 or more AML cell lines are shown. Side box plots demonstrate expression distribution in AML cell lines (Broad CCLE database) and primary AML samples.

**Supplementary Fig. 2. Prediction of MHC-II gene expression by CORENODE. a,** Combinatorial regression of MHC-II genes. Comparison of leave-one-out error (LOO) and *r*^2^ for 1-, 4- and 5-mers describing expression of 6 MHC type II genes (HLA-DPA1, HLA-DBP1, HLA-DQA1, HLA-DQB1, HLA-DRA, HLA-DRB1). 5-mers where CIITA is the 5^th^ term display the smallest LOO and the largest *r*^2^ values indicating improved fit without overfitting. **b**, Gene expression in two populations of the TCGA dataset. The blue population represents patient samples with below-median expression of IRF8 and MEF2C and above-median expression of MYB and MEIS1, while the red population represents patient samples with the opposite pattern of TF expression.

**Supplementary Fig. 3. Cell surface expression of MHC-II genes following TF knockout. a**, Validation of gene knockouts. MV411 cells were electroporated with RNPs targeting the indicated genes and protein depletion was verified by Western blot. Unless indicated otherwise, imaging was performed 72 hours post electroporation. **b**, FACS was performed using a pan-specific anti-HLA-DP/DQ/DR antibody in the indicated AML cell lines at indicated times after TF knockout. Data represent an average of 3 replicates and are normalized to the AAVS1 (“safe harbor” targeting) sgRNA control. Asterisks represent *p*-value <0.05 calculated by the two-tailed *t*-test.

**Supplementary Fig. 4. Histone and TF architecture in the MHC-II locus. a,** H3K27ac metatracks. *Red*, tracks from 10 AML patients with the highest MHC type II expression; *blue*, tracks from the 10 AML patients with the lowest MHC type II expression. Each metatrack is a collection of semi-transparent area plots representing individual samples and the average profile is represented by a thick line. **b**, Heatmap of aggregated SE activities associated with the MHC-II locus and the tetrad TFs. H3K27ac ChIP-seq data are from Ref.(*9*). Multiple SEs were called by ROSE2 for each locus and their signal was aggregated and z-scored. Rows and columns were hierarchically clustered by Pearson correlation with complete linkage. **c**, H3K27 acetylation and TF binding in the MHC type II locus. Green tracks are H3K27ac metatracks composed by overlaying semi-transparent area plots representing 2 biological replicates of the indicated TF knockouts compared to a control locus-targeting sgRNA (AAVS1). The average profile is represented by a thick line. Grey tracks represent binding of the indicated TFs. All of the shown ChIP-seq experiments were performed in MV411 except for the CIITA track which was downloaded from Ref.(*29*) and represents a lymphoid cell line. **d**, Density plots of spike-in controlled H3K27ac ChIP-seq experiment showing genomewide histone acetylation changes after indicated TF knockouts, compared to a control locus-targeting sgRNA (AAVS1). Each row represents a single peak.

**Supplementary Fig. 5. CIITA and a GvL response module. a**, Correlation between CIITA and MHC-II gene expression in the BeatAML dataset. Note that these plots represent Pearson correlation and are different from CIITA 1-mers (such as the one in Fig. 2A) which have a quadratic term in addition to a linear term. **b**, Validation of a CIITA knockout in MV411 cells by PCR amplification followed by agarose gel electrophoresis with ethidium bromide staining. Three closely positioned gRNAs were used for the CIITA knockout, producing a size shift on gel electrophoresis indicating complete editing efficiency in the bulk population and a homogenous excision in the clonally selected line. **c**, A genome-wide heatmap and clustering of edges. Edges with scores ≥8 were converted to 1 for predicted positive regulation (DD > 0) or −1 for negative regulation (DD < 0). The resulting matrix was clustered by Pearson correlation (CR TFs (columns): hierarchical, complete linkage; target genes (rows): k-means with k=15) using Morpheus (https://software.broadinstitute.org/morpheus). **d**, Gene-set enrichment analysis of the GvL module. The ChEA dataset shows enrichment of TF binding in the gene promoters, while the KEGG dataset shows gene pathway enrichment. Enrichments were computed using the Enrichr (*30*). **e**, Expression of the GvL module in paired primary and post-chemotherapy relapse patient samples from Ref.(*3*).

**Supplementary Fig. 6. MHC-II expression and myeloid differentiation status. a**, Expression of the 19 myeloid markers in sorted normal progenitors. **b**, Normal progenitors and leukemic cells are plotted according to their indices of myeloid and lymphoid development. Panels **a** and **b** were computed with data from Ref.(*31*). **c** and **d**, Poor correlation MHC-II expression and myeloid differentiation state. A composite score reflecting MHC-II expression (HLA index) is plotted against the index of myeloid development using BeatAML (*15*) and TCGA (*16*) datasets, respectively.

## METHODS

### External datasets

RNA-seq BAM files from the BeatAML project (*15*) were provided by Oregon Health & Science University and processed through the CCLE RNA processing pipeline (STAR/RSEM, described at https://github.com/broadinstitute/ccle_processing). Reads were normalized to transcripts per million (TPM) and filtered for protein coding genes. The expression values were transformed to log_2_(TPM+1). The genes were ranked by average expression across the samples and 9000 top expressed genes were chosen for further analysis (Supplementary Fig. 1).

H3K27ac ChIP-seq data from primary AML samples (*9*) were downloaded from Sequence Read Archive (SRA) under accession number SRP103200 and processed using the AQUAS pipeline (https://github.com/kundajelab/chipseq_pipeline) with minor modifications and according to the ENCODE3 guidelines. We used data from 49 samples that passed our quality criteria. Reads were aligned to the hg19 genome build using BWA-ALN and peaks were called using MACS2. For superenhancer (SE) calling each sample was processed using ROSE2 (https://github.com/linlabbcm/rose2), excluding 2500 bp around TSSs (-t 2500) and the hg19 Encode blacklisted regions. SE regions were then merged and ROSE2 was run again for each sample on the merged regions, producing the SE signal matrix, which was then normalized by median signal for each sample.

Paired processed RNA-seq data from the two available studies of AML patients at the time of initial presentation and relapse (*3, 4*) were downloaded from https://www.nejm.org/doi/full/10.1056/NEJMoa1808777 and https://www.ebi.ac.uk/arrayexpress/experiments/E-MTAB-7456/samples/.

Genetic dependency data are available for download at the Broad DepMap portal database (https://depmap.org/portal/download/). Data release 20q1 was used for this study. For the purpose of this study we considered a gene to represent a selective AML dependency if it had a probability of dependency of ≥0.5 in 3 or more AML cell lines and not be considered a pan-lethal dependency based on the LRT score. This approach resulted in a total of 225 selective AML dependencies.

### Cell culture

AML cell lines were purchased from the American Type Culture Collection (https://www.atcc.org). Cells were cultured in the RPMI-1640 media containing 10% fetal bovine serum and regularly tested to be free of *Mycoplasma spp*.

### CRISPR-Cas9 gene knockouts

Synthetic modified sgRNA constructs were purchased from Synthego (Redwood City, CA; refer to **Error! Reference source not found.** for sgRNA sequences). Ribonucleoprotein (RNP) assembly was performed by mixing 2-3 sgRNAs (a total of 120 pmol) with 8.5 μg recombinant Cas9 (Invitrogen A36499). The resulting RNP mix was electroporated into 0.3E6 MV411 cells using a Lonza 4D Nucleofector, program DJ-100, in 20 μl Nucleocuvette strips (Lonza V4XC-2032). Unless otherwise noted, cells were incubated in media for 72 hours post-electroporation before subsequent analyses. Knockout efficiency was confirmed by Western blotting and PCR amplification followed by indel analysis. A guide RNA targeting the AAVS1 “safe harbor” locus was used as a negative control (*32*).

### Genome-wide occupancy analysis

ChIP-seq was performed as previously described (*33*) and in accordance with the Encode guidelines (*34*). The following antibodies were used for ChIP: H3K27ac (Abcam ab4729), IRF8 (SantaCruz sc365042X), MYB (Abcam ab45150), MEIS1 (Abcam ab19867) and MEF2C (GeneTex GTX105433). ChIPseq libraries were prepared using Swift S2 Acel reagents (Swift 21096) on a Beckman Coulter Biomek i7 liquid handling platform from approximately 1 ng of DNA according to manufacturer’s protocol and using 14 cycles of PCR amplification. Sequencing libraries were quantified by Qubit fluorometer and Agilent TapeStation 2200. Library pooling and indexing was evaluated by shallow sequencing on Illumina MiSeq. Subsequently, libraries were sequenced on NovaSeq targeting 40 million 100 bp read pairs by the Molecular Biology Core facilities at the Dana-Farber Cancer Institute.

For quantitative ChIP-seq analysis of H3K27 acetylation we used *Drosophila* chromatin/antibody spike-in control as previously described (*35*). Briefly, 4 μg of anti-H3K27ac antibody, 2 μg of spike-in antibody and 20 ng of spike-in chromatin (Active Motif 61686 and 53083, respectively) were added to chromatin prepared from 2.5×10^6^ MV411 cells 72 hours after RNP-mediated TF knockout. The rest of the ChIP-seq experiment was performed in the standard fashion. After ChIP-seq reads were mapped to the *Drosophila* genome and the hg38 human genome in parallel, human tag counts were normalized to *Drosophila* tag counts.

### Global transcriptome analysis

For RNA-seq experiments the total cellular RNA was extracted using the QuickRNA kit (Zymo Research R1054). Purified total RNA was mixed with the ERCC synthetic spike-in control (Invitrogen 4456740). RNA sequencing libraries were prepared on a Beckman Coulter Biomek i7 liquid handling platform using Roche Kapa mRNA HyperPrep strand specific sample preparation kits (Roche 08098123702) from 200 ng of purified total RNA according to the manufacturer’s protocol. Library quantification and Illumina sequencing were performed as described in the ChIP-seq section above.

### Measurement of MHC-II expression by FACS

For MHC Class II analysis, cells were stained with a pan-specific FITC-conjugated Mouse Anti-Human HLA-DR, DP, DQ antibody (BD Biosciences 555558, clone Tu39), and a PE-conjugated Mouse Anti-Human HLA-DR antibody (BD Biosciences 555812, clone G46-6). Data were analyzed in FlowJo (FlowJo LLC, Ashland, OR).

### Western blotting

Whole-cell lysates were prepared in RIPA buffer (Boston Bio-Products BP-115-500) with protease inhibitor cocktail (ThermoFisher 23225). Lysates were boiled in Laemmli buffer (BioRad 1610737), separated by SDS-PAGE, and transferred and blocked using standard methodology. HRP-conjugated anti-mouse and anti-rabbit IgG secondary antibodies were used for imaging (BioRad 1706515 and 1706515) with an enhanced chemiluminescence substrate (PerkinElmer NEL104001EA) according to manufacturers’ instructions. The following primary antibodies were used: anti-MYB (Abcam ab45150), anti-MEF2C (Cell Signaling 5030S), anti-MEIS1 (Abcam ab19867), anti-IRF8 (Cell Signaling 5628S).

### Statistical analysis

All experiments were done in at least 3 replicates. Two-tailed Student’s *t* test was used to compare mean fluorescence of antibody-stained cells following TF knockout. Changes in gene expression in patient samples between initial diagnosis and relapse were ascertained using a Wilcoxon signed-rank test. Statistical analysis of ChIP-seq and RNA-seq data was performed using the DEseq2 package(*36*).

### Myeloid differentiation index

To identify markers of myeloid development, genome-wide mRNA expression values in “HSC” and “monocyte” samples from Ref.(*31*) were processed to yield mean and variance of expression by gene. For each gene, the two variances were pooled [pooled variance = mean (HSC variance, monocyte variance)]. A separation index was then defined for each gene as the difference between the HSC and monocyte mean expressions divided by the square root of the pooled variance. Markers were chosen as the 19 genes with the highest separation indices. Lymphoid markers were identified with the same procedure, comparing HSC samples to the T-cell and B-cell samples. To compute the myeloid index, each sample’s expression of the marker genes as determined above was converted to a z-score using the mean of all eight HSC and monocyte expressions and the pooled variance for that gene, and further normalized to a +/- 1 scale by dividing by the maximum absolute value of all z-scores. For genes where the mean monocyte score was higher than the mean HSC score, normalized scores were multiplied by (−1). The normalized scores for the set of marker genes were summed to define the myeloid index for each sample. The same procedure was used for the lymphoid index, with appropriate cell-type substitution. Plotting normal and leukemia cell types according the myeloid and lymphoid indices yielded the expected development vectors as shown in Supplementary Fig. 6. The same markers were then used to define a myeloid index for the samples from the other datasets (Refs.(*3, 4, 15, 16*)). The HLA index was defined as the mean expression (in log2[TPM+1]) of the nine HLA-D genes.

## Acknowledgements

M. Pimkin is supported by a Damon Runyon-Sohn Pediatric Cancer Fellowship and a Young Investigator Award from the Alex’s Lemonade Stand Foundation. This work was supported, in whole or in part, by research grants to M. Pimkin from the Curing Kids Cancer Foundation, When Everyone Survives Foundation, Pedals for Pediatrics, Children’s Cancer Research Fund, Children’s Leukemia Research Association, Hyundai Hope on Wheels, Kate Amato Foundation, and Boston Children’s Hospital Office of Faculty Development. K Stegmaier was supported by NIH 5R35 CA210030 and NIH P50 CA206963. N.V. Dharia was supported by the Julia’s Legacy of Hope St. Baldrick’s Foundation Fellowship. We thank Drs. L.S. Kean, G.C. Yuan and S. Avagyan for critical reading of this manuscript, and Drs. T. Ley and C. Miller for access to the scRNA-seq data.

## Author contributions

Conceptualization: MP, SHO, KE, VRP, CYL

Computational methodology: KE, MPi, JK, MPe, JVK, JMD, GK, NVK

Experiments: TH, MP, YH, JE

Visualization: MPe, KE, MPi

Funding acquisition: SHO, MPi

Supervision: MPi, SHO, KS

Writing and review of the manuscript: all authors

