## Supplementary Figures for "Transcriptional plasticity drives leukemia immune escape"

### Supplementary Figure 1

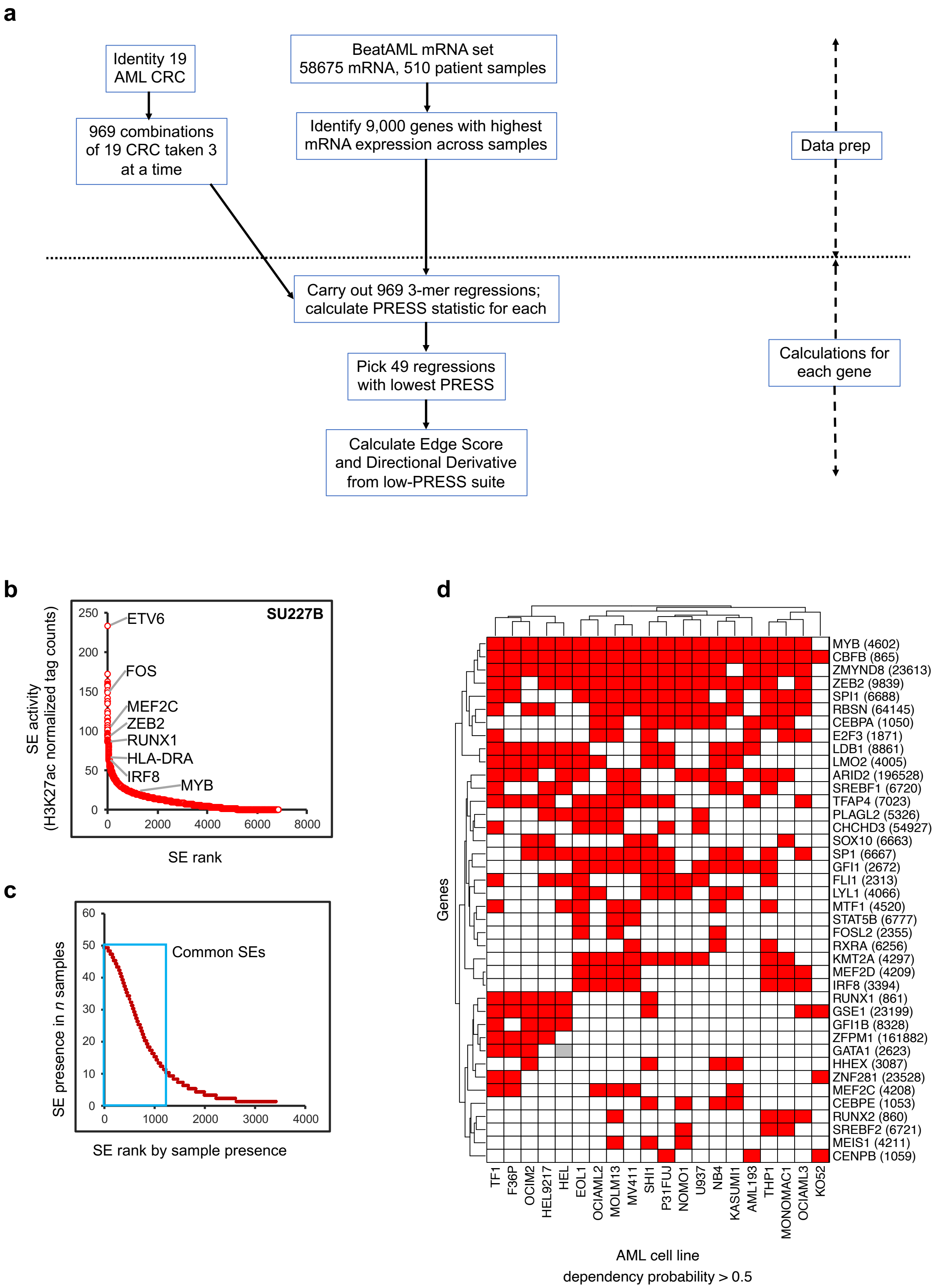

### Supplementary Figure 2

**a**

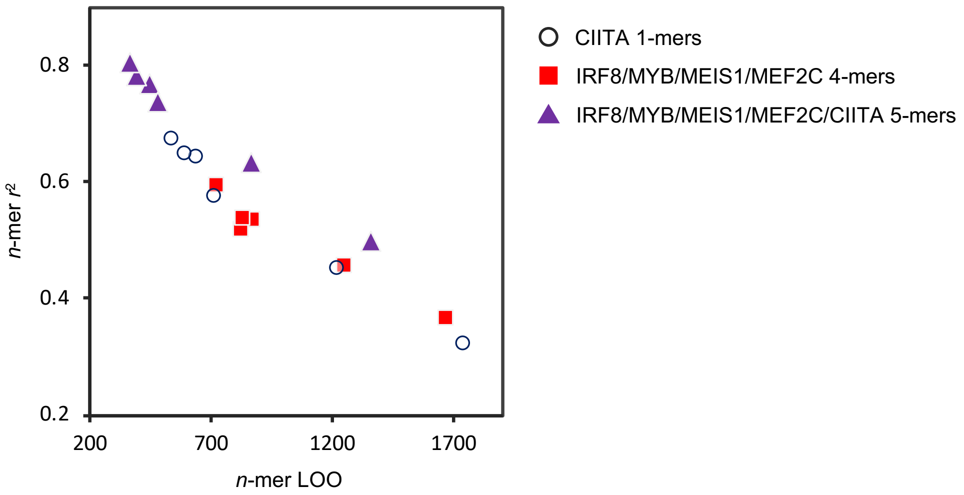


**b**

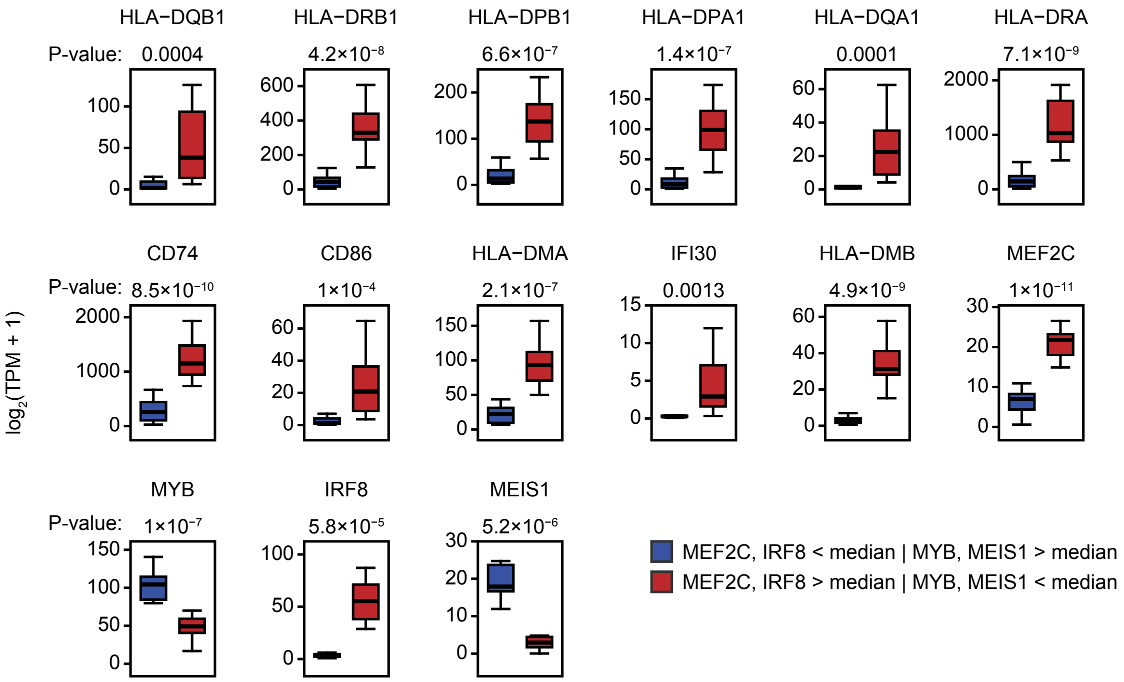

#### Supplementary Figure 4

**a**

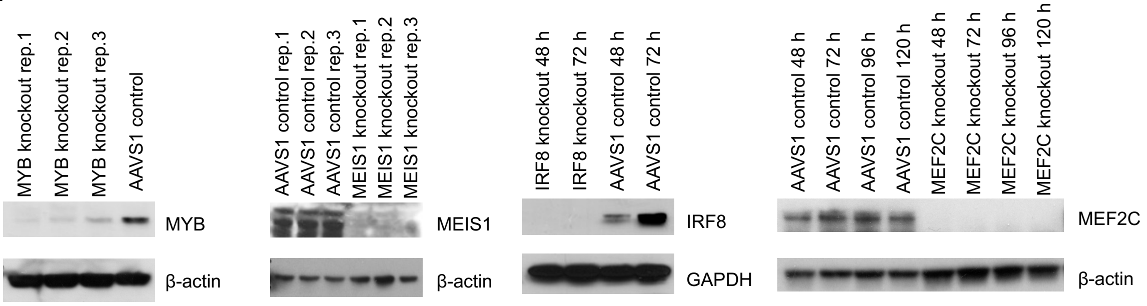


**b**

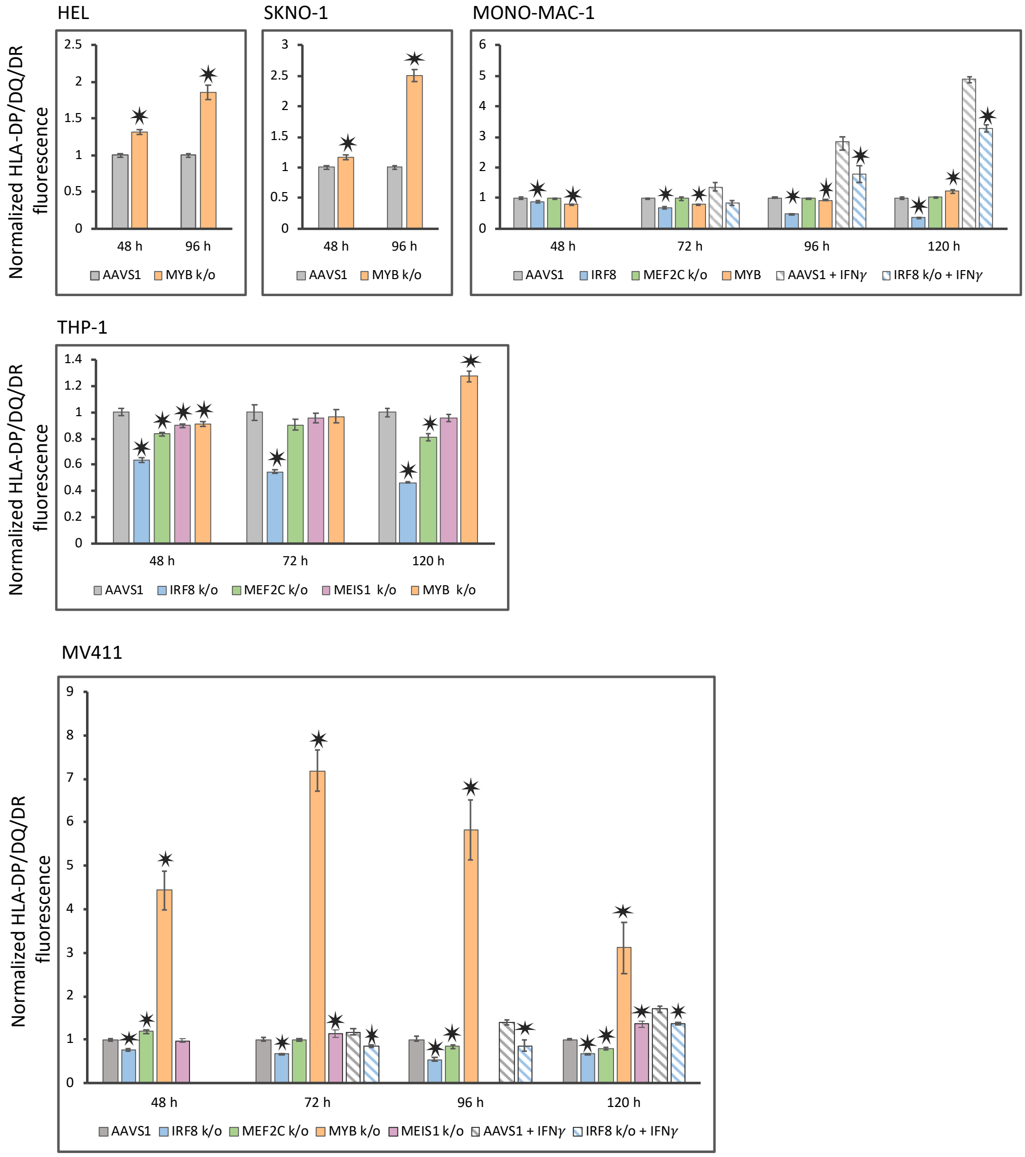

### Supplementary Figure 4

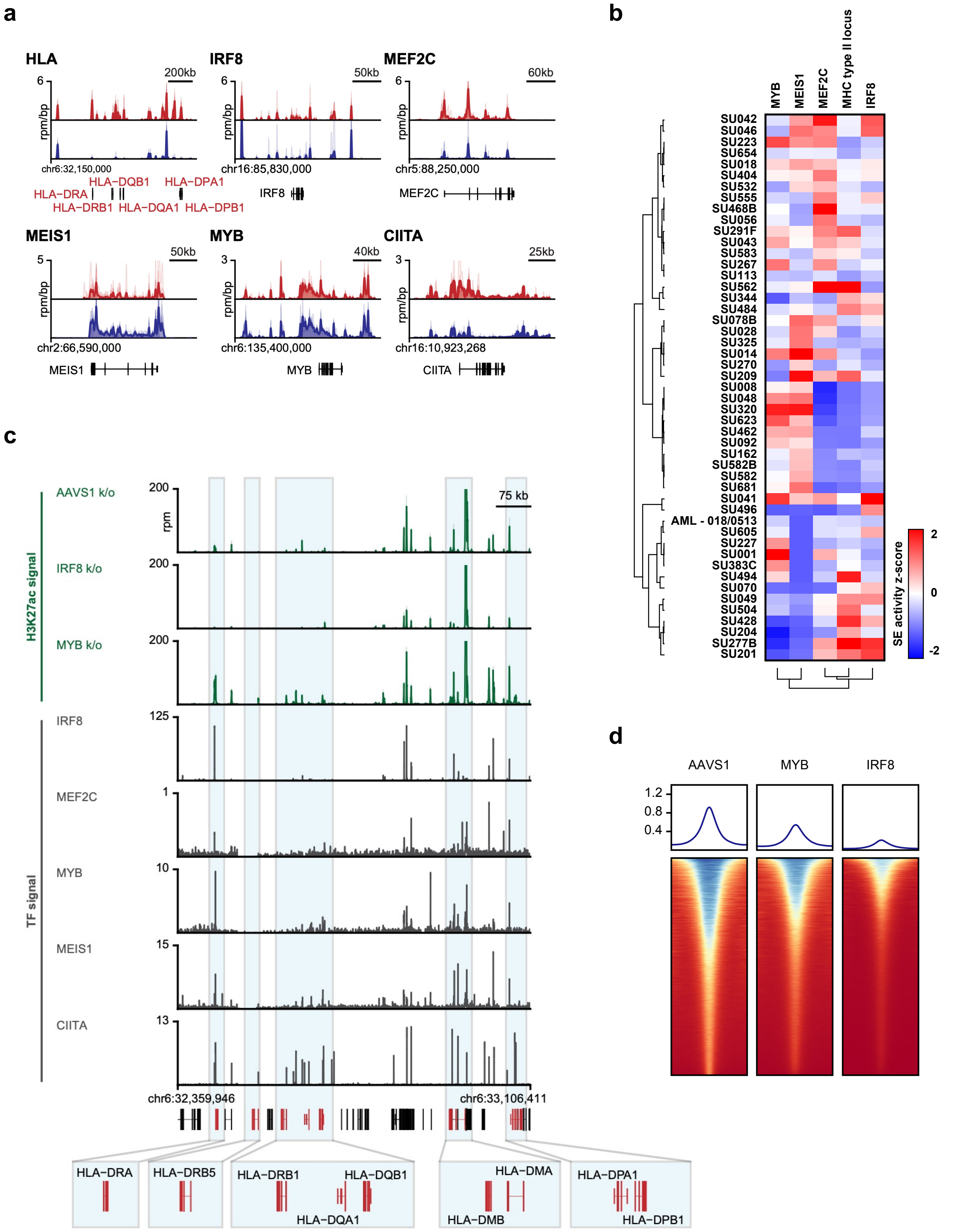

### Supplementary Figure 5

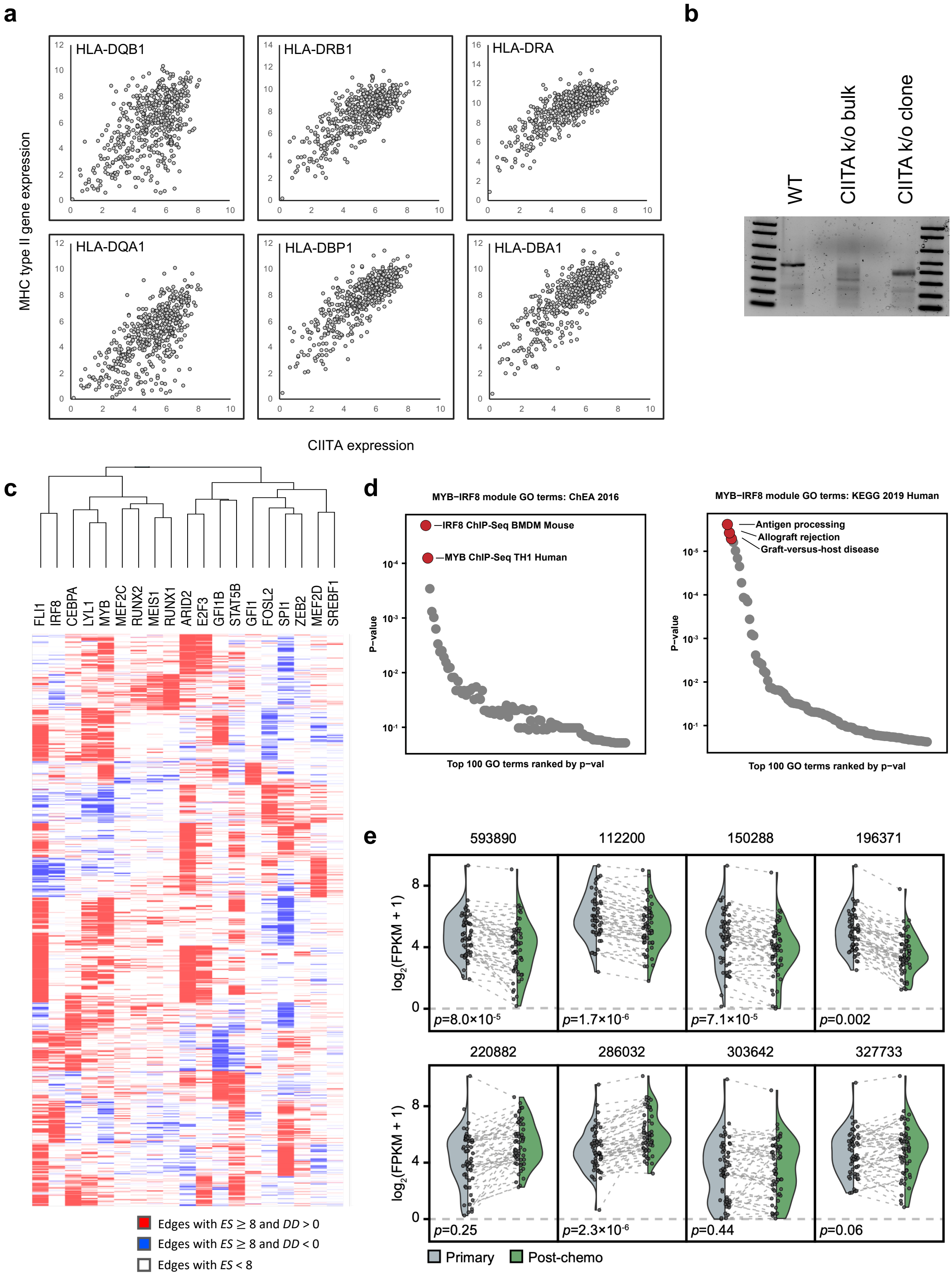

### Supplementary Figure 6

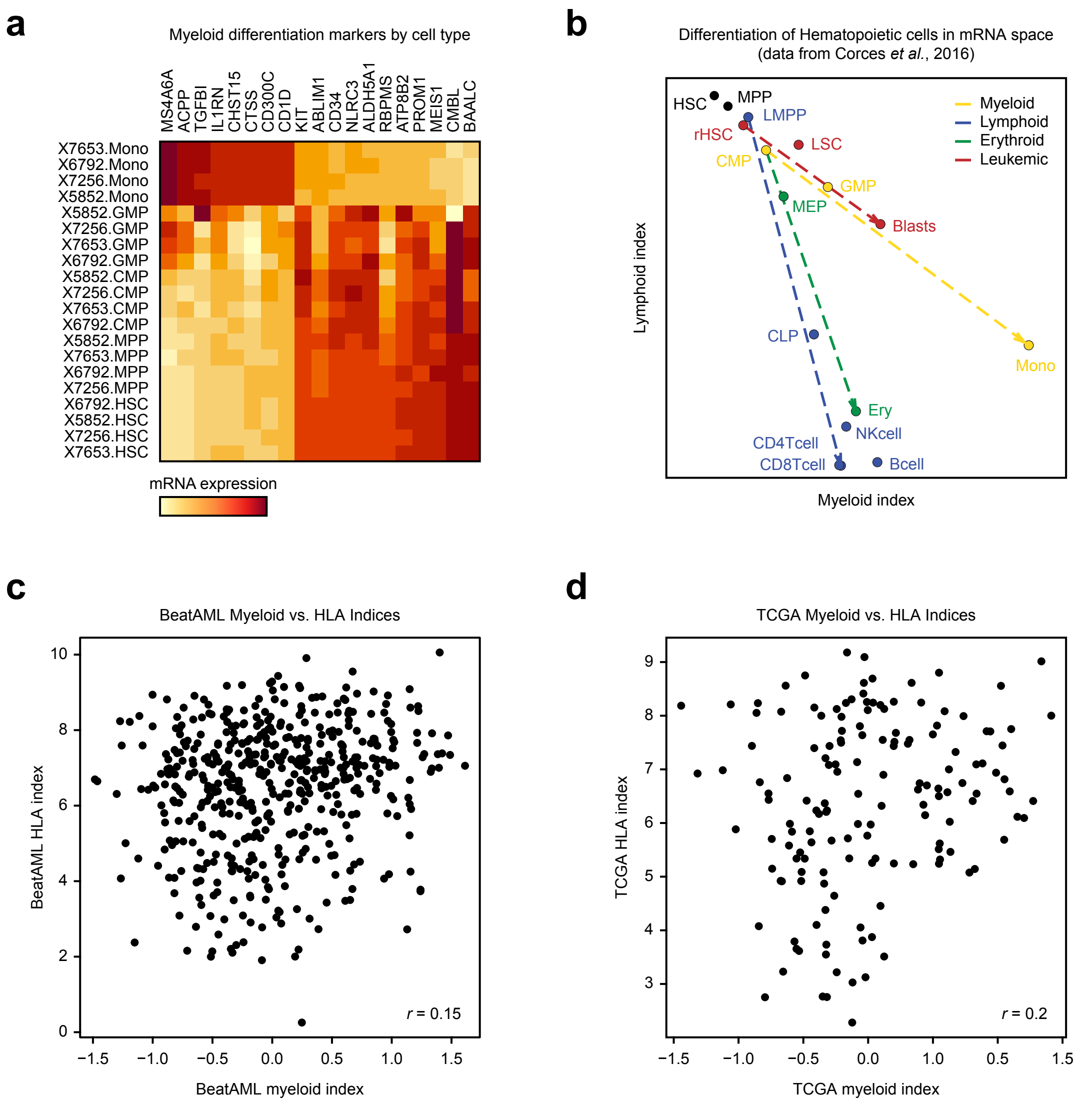
